# AStarix: Fast and Optimal Sequence-to-Graph Alignment

**DOI:** 10.1101/2020.01.22.915496

**Authors:** Pesho Ivanov, Benjamin Bichsel, Harun Mustafa, André Kahles, Gunnar Rätsch, Martin Vechev

## Abstract

We present an algorithm for the *optimal alignment* of sequences to *genome graphs*. It works by phrasing the edit distance minimization task as finding a shortest path on an implicit alignment graph. To find a shortest path, we instantiate the A^⋆^ paradigm with a novel domain-specific heuristic function that accounts for the upcoming subsequence in the query to be aligned, resulting in a provably optimal alignment algorithm called AStarix.

Experimental evaluation of AStarix shows that it is 1–2 orders of magnitude faster than state-of-the-art optimal algorithms on the task of aligning Illumina reads to reference genome graphs. Implementations and evaluations are available at https://github.com/eth-sri/astarix.

## 1 Introduction

The analysis and understanding of genetic variation encoded in the genome of an organism lies at the center of computational biology and medicine. Variation is usually identified through matching sequences obtained from DNA/RNAsequencing back to a reference (genome) sequence in the process of *variant calling*, making the alignment task a core problem in sequence bioinformatics.

Historically, a single linear reference sequence has been used to represent the most common variants in a population. While providing a working abstraction for most cases, rare or sub-population specific variation is especially hard to model in this setting, creating a reference allele bias [35,4]. Consequently, in the last few years, the field has shifted first towards using sets of reference sequences, and more recently to graph data structures (so-called *genome graphs*), to represent many genomes or haplotypes simultaneously [7,25,9].

Both for sequence-to-sequence alignment and sequence-to-graph alignment, heuristics are employed to keep alignment tractable [2,21,9], especially for large populations of human-sized genomes. While such heuristics find the correct alignment for simple references, they often perform poorly in regions of very high complexity, such as in the human major histocompatibility complex (MHC) [7], in complex but rare genotypes arising from somatic-subclones in tumor sequencing data [10], or in the presence of frequent sequencing errors [29]. Importantly, these cases can be of specific clinical or biological interest, and incorrect alignment can cause severe biases for downstream analyses. For instance, the combination of high variability of MHC sequences in humans and small differences between alleles [5] leads to a risk of misclassifications due to suboptimal alignment. Guaranteeing optimal alignment against all variations represented in a graph is a major step towards alleviating those biases.

Formally, we consider the optimal *sequence-to-graph alignment* problem, the task of finding an optimal base-to-base correspondence between a query sequence and a (possibly cyclic) walk in the graph. Related alignment problems have already been formulated as graph shortest path problems [3,16].

### 1.1 Related Work

#### Seed-and-Extend

Since optimal alignment is often intractable, many aligners use heuristics, most commonly the *seed-and-extend* paradigm [2,21,22]. In this approach, alignment initiation sites (*seeds*) are determined, which are then *extended* to form the *alignments* of the query sequence. The fundamental issue with this approach, however, is that the seeding and extension phases are mostly decoupled during alignment. Thus, an algorithm with a provably optimal extension phase may not result in optimal alignments due to the selection of a suboptimal seed in the first phase. In cases of high sequence variability, the seeding phase may even fail to find an appropriate seed from which to extend.

#### Accounting for Variation

First attempts to include variation into the reference data structure were made by augmenting the local alignment method to consider alternative walks during the extend step [30,17]. This approach has since been extended from the linear reference case to graph references. To represent non-reference variation of multiple references during the seeding stage, HISAT2 uses generalized compressed suffix arrays [33] to index walks in an augmented reference sequence, forming a local genome graph [19]. VG [9] uses a similar technique [32] to index variation graphs representing a population of references.

BrownieAligner, another recent work developed for local alignment of sequences to *de Bruijn* graph representations of genomic variation, features an optimal extension phase using a branch-and-bound-based early cutoff, while employing a heuristic maximal-exact-match approach for seeding [11].

#### Optimal Alignment

Current optimal sequence-to-graph alignment algorithms reach their worst-case 𝒪(*nm*) runtime [16]. In this light, approaches for improving the efficiency of optimal alignment have taken advantage of specialized features of modern CPUs to improve the practical runtime of the Smith-Waterman dynamic programming (DP) algorithm [34] considering all possible starting nodes. These use modern SIMD instructions (e.g. VG [9] and PaS-GAL [15]) or reformulations of edit distance computation to allow for bit-parallel computations in GraphAligner^1^ [27]. Many of these, however, are designed only for specific types of genome graphs, such as *de Bruijn* graphs [24,11,23] and variation graphs [9]. A compromise often made when aligning sequences to cyclic graphs using algorithms reliant on directed acyclic graphs involves the computationally expensive “DAG-ification” of graph regions [18,9].

#### A^⋆^ algorithm

We aim to guarantee optimal alignment while optimizing the average runtime to not reach its worst case complexity. While Dijkstra is an algorithm that explores graph nodes in the order of their distance from the start, A^⋆^ is a generalization of Dijkstra that also accounts for their distance from the target. A^⋆^ prioritizes the exploration of nodes that seem to be closer to the target nodes. This way, A^⋆^ can sometimes dramatically improve on the performance of Dijkstra while remaining optimal.

There has been one attempt to apply A^⋆^ for optimal alignment [8] which uses a heuristic function that accounts only for the length of the remaining query sequence to be aligned. However, it does not significantly outperform Dijkstra (in fact, it is equivalent for a zero matching cost). In contrast, the heuristic function we introduce is more informative and consistently outperforms Dijkstra.

### 1.2 Main Contributions

We introduce a novel approach, called AStarix, for optimal sequence-to-graph alignment based on A^⋆^. As with any A^⋆^ instantiation, the core difficulty lies in developing an accurate domain-specific heuristic which is fast to compute. We design a heuristic that accounts for the content of the upcoming query letters to be aligned, which more effectively guides the search. Our proposed heuristic has two advantages: (i) it is correctness-preserving, that is, it preserves the fact that AStarix finds the best alignment, yet (ii) it is practically effective in that the algorithm performs a near-optimal number of steps. Overall, this heuristic enables AStarix to compute the best alignment while also scaling to larger reference graph sizes when compared to existing state-of-the-art optimal aligners.

Our main contributions^2^ include:

#### 1. AStarix

An algorithm for optimal sequence-to-graph alignment based on a novel instantiation of A^⋆^ with an accurate domain-specific heuristic that accounts for the upcoming query letters to be aligned (§3).

#### 2. Algorithmic optimizations

To ensure that AStarix is practical, we introduce a number of algorithmic optimizations which increase performance and decrease memory footprint (§4). We also prove that all optimizations are correctness-preserving.

#### 3. Thorough experimental evaluation of AStarix

We demonstrate that AStarix is up to 2 orders of magnitude faster than other optimal aligners on various reference graphs (§5).

## 2 Task Description: Alignment to Reference Graphs

We analogously to reference graphs, now describe the task of aligning a query to a reference graph. To this end, we (i) introduce the task of optimal alignment on a *reference graph*, (ii) formalize this task in terms of an *edit graph*, and (iii) introduce an alternative formulation in terms of an *alignment graph*, which is the basis for shortest path formulations of the optimal alignment. Fig. 1 summarizes these different graph types.

**Fig. 1:**
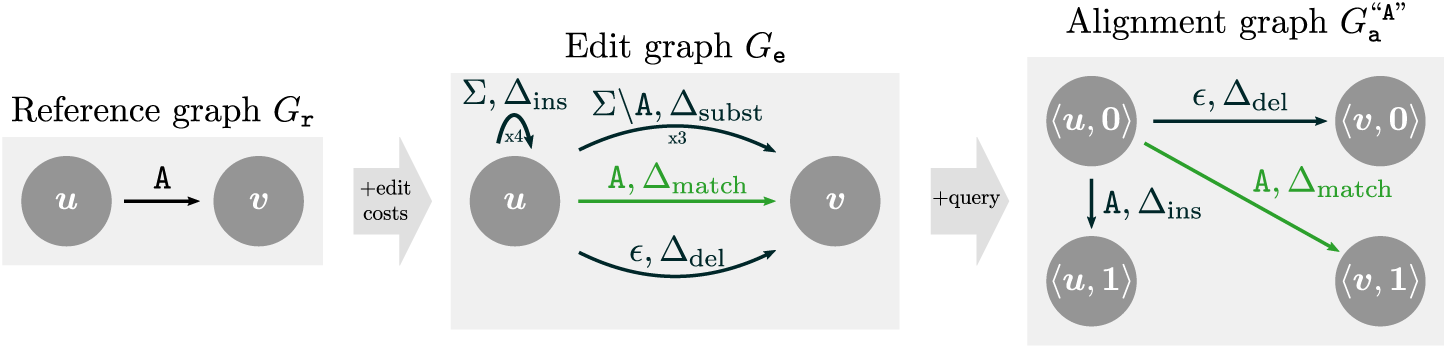
Starting from the reference graph (left), we can construct the edit graph (middle) and the alignment graph 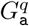 for query *q* = “`A`” (right). Edges are annotated with labels and/or costs, where sets of labels represent multiple edges, one for each letter in the set (indicated by “x3” and “x4”).

### Reference Graph

We encode the collection of references to which we want to align in a reference graph, which captures genomic variation that a linear reference cannot express [25,9]. We formalize a reference graph as a tuple *G*_r_ = (*V*_r_, *E*_r_) of nodes *V*_r_ and directed, labeled edges *E*_r_ ⊆*V*_r_ ×*V*_r_ × *Σ*, where the alphabet *Σ* = {`A`, `C`, `G`,}`T` represents the four different nucleotides. Note that in contrast to sequence graphs [28], we label edges instead of nodes.

### Path, Spelling

Any path *π* = (*e*_1_, …, *e*_*k*_) in *G*_r_ induces a *spelling σ* (*π*) ∈ *Σ*^*^ defined by *σ*(*e*_1_) *…σ*(*e*_*k*_), where *σ*(*e*_*i*_) is the label of edge *e*_*i*_ and *Σ*\* ≔ ∪_*k*∈ℕ_ *Σ*^*k*^ We note that our approach naturally handles cyclic walks and does not require cycle unrolling, a feature shared with BitParallel [27] and BrownieAligner [11] but missing from VG [9], PaSGAL [15] and V-ALIGN [18].

### Alignment on Reference Graph

An *alignment* of *query q*∈ *Σ*^*^ to a reference graph *G*_r_ = (*V*_r_, *E*_r_) consists of (i) a path *π* in *G*_r_ and (ii) a sequence of edit operations (matches, substitutions, insertions, deletions) transforming *σ*(*π*) to *q*.

### Optimal Alignment, Edit Distance

Each edit operation is associated with a real-valued cost (Δ_match_, Δ_subst_, Δ_ins_, and Δ_del_, respectively). An optimal alignment minimizes the total cost of the edit operations converting *σ*(*π*) to *q*. For optimal alignments, this total cost is equal to the edit distance between *σ*(*π*) and *q*, i.e., the cheapest sequence of edit operations transforming *σ*(*π*) into *q*.

We make the (standard) assumption that 0 ≤Δ_match_ ≤Δ_subst_, Δ_ins_, Δ_del_, which will be a prerequisite for the correctness of our approach.

### Edit Graph

Instead of representing alignments as pairs of (i) paths in the reference graph and (ii) sequences of edit operations on these paths, we introduce *edit graphs* whose paths intrinsically capture both. This way, we can formally define an alignment more conveniently as a path in an edit graph.

Formally, an *edit graph G*_e_ := (*V*_e_, *E*_e_) has directed, labeled edges *E*_e_ ⊆ *V*_e_ × *V*_e_ × *Σ*_*ϵ*_ × ℝ_≥0_ with associated costs that account for edits. Here, *Σ* _∈_:= *Σ* ∪{∈} extends the alphabet *Σ* by ∈ to account for deleted characters (see Fig. 1). The edit and reference graphs consist of the same vertices, i.e., *V*_e_ = *V*_r_. However, *E*_e_ contains more edges than *E*_r_ to account for edits. Concretely, for each edge (*u, v, l*) ∈*E*_r_, *E*_e_ contains edges to account for (i) matches, by an edge (*u, v, l*, Δ_match_), (ii) substitutions, by edges (*u, v, l*′, Δ_subst_) for each *l*′ *Σ l*, (iii) deletions, by an edge (*u, v, E*, Δ_del_), and (iv) insertions, by edges (*u, u, l*′, Δ_ins_) for each *l*′∈ *Σ*. The spelling *σ*(*π*) *Σ*^*^ of a path *π*; ∈ *G*_e_ is defined analogously to reference graphs, except that deleted letters (represented by *E*) are ignored. The cost cost(*π*) of a path *π* ∈ *G*_e_ is the sum of all its edge costs.

### Alignment on Edit Graph

An *alignment* of query *q* to *G*_r_ is a path *π* in *G*_e_ spelling *q*, i.e., *q* = *σ*(*π*). An *optimal alignment* is an alignment of minimal cost

### Alignment Graph

To find an optimal alignment of *q* to the edit graph *G*_e_ using shortest path finding algorithms, we must ensure that only paths spelling *q* are considered. To this end, we introduce an alternative but equivalent formulation of alignments in terms of an *alignment graph* 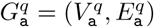.

Here, each *state* 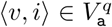 consists of a vertex *v* ∈ *V*_e_ and a query position *i* ∈ {0, …, |*q*|} (equivalent to [28]). Traversing a state 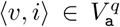. Represents the alignment of the first *i* query characters ending at node *v*. In particular, query position *i* = 0 indicates that we have not yet matched any letters from the query. We note that the alignment graph explicitly depends on the query *q*. In particular, the example alignment graph 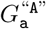. in Fig. 1 lacks substitution edges from *G*_e_, as their labels (`C`, `G`, `T`) do not match the query *q* = “*A*”.

We construct the alignment graph 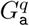.to guarantee that any walk from a source ⟨*u*, 0⟩ to a state ⟨*v, i*⟩ corresponds to an alignment of the first *i* letters of query *q* to *G*_r_. As a consequence, there is a one-to-one correspondence between alignments *π*_e_ of *q* to *G*_e_ and paths 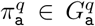 from sources *S* := *V*_r_ × {0} to targets *T* := *V*_r_ × {|*q* |}, with 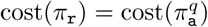. To find the best alignment in *G*_e_, only paths in 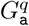. (walks without repeating nodes) can be considered, since repeating a node in 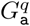. cannot lead to a lower cost (Δ_del_ ≥ 0) for the same state.

The edges 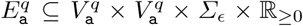.are built based on the edges in *E*_e_, except that the former (i) keep track of the position in the query *i*, and (ii) only contain empty edges or edges whose label matches the next query letter:

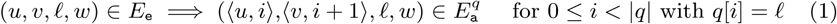

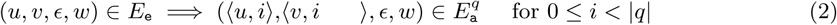

Here, assuming 0-indexing, *q*[*i*] is the next letter to be matched after matching *i* letters. Then, Eq. (1) represents matches, substitutions, and insertions (which advance the position in the query by 1), while Eq. (2) represents deletions (which do not advance the position in the query).

#### Algorithm 1 AStarix including heuristic function.

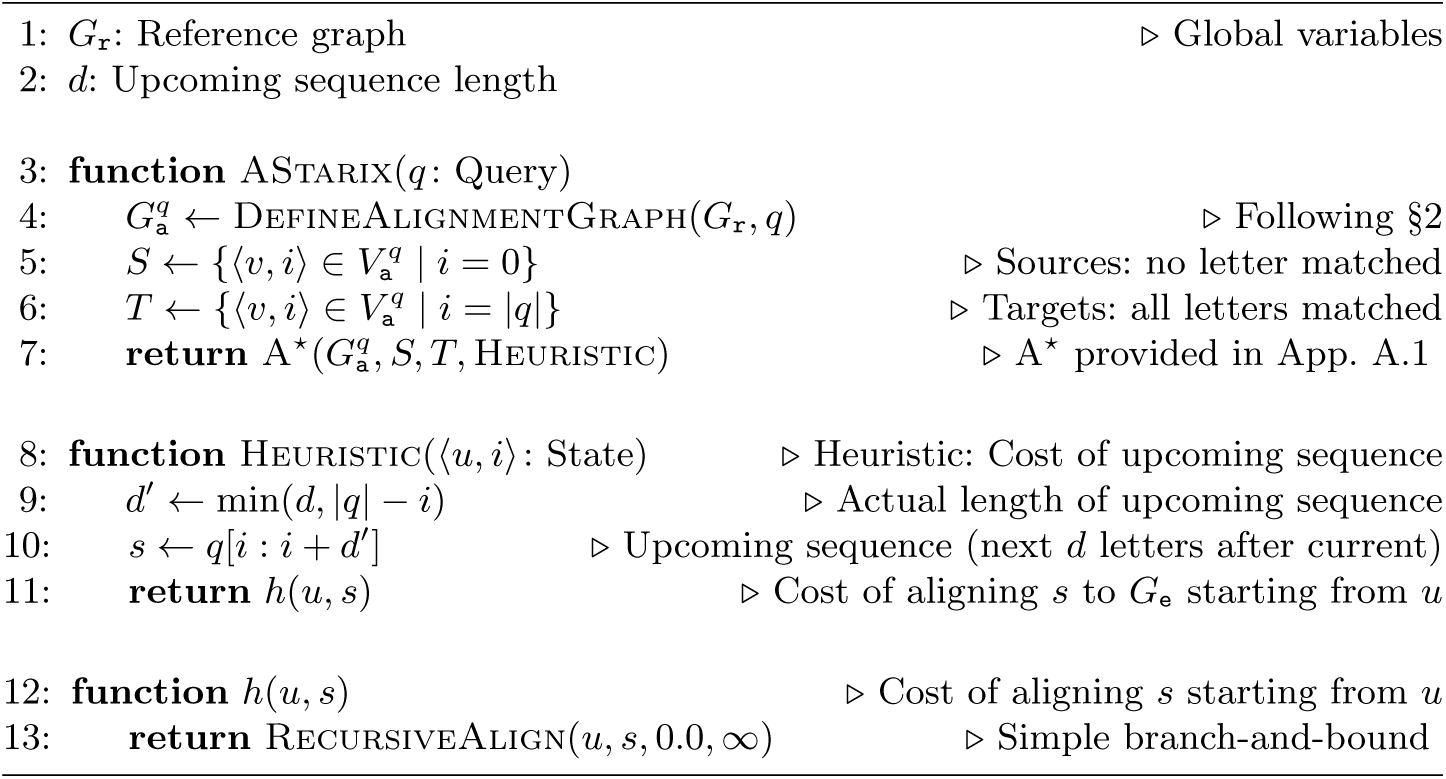

### Dynamic Construction

As the size of the alignment graph is 𝒪(|*G*_r_| |*q*|), it is expensive to build it fully for every new query. Therefore, our implementation constructs the alignment graph 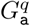 on-the-fly: the outgoing edges of a node are only generated on demand and are freed from memory after alignment.

## 3 AStarix: Finding Optimal Alignments Using A^⋆^

In this section, we first introduce the general A^⋆^ algorithm for finding shortest paths, and the notion of an optimistic heuristic, a sufficient condition for instantiations of A^⋆^ to be correct (i.e., to indeed find shortest paths). Then we instantiate A^⋆^ with our domain-specific heuristic that accounts for upcoming subsequences to be aligned, and prove that this heuristic is optimistic.

### 3.1 Background: General A^⋆^ algorithm

Given a weighted graph *G* = (*V, E*) with *E* ⊆ *V* × *V* × ℝ_≥0_, the A^⋆^ algorithm (abbreviated as A^*^) searches for the shortest path from sources *S* ⊆ *V* to targets *T* ⊆ *V*. It is an extension of Dijkstra’s algorithm that additionally leverages a *heuristic function h* : *V* →ℝ_≥0_ to decide which paths to explore first. If *h*(*u*) ≡ 0, A^⋆^ is equivalent to Dijkstra’s algorithm. We provide an implementation of A^⋆^ and Dijkstra in App. A.1, but do not assume knowledge of either algorithm in the following. At a high level, A^⋆^ maintains the set of all *explored* states, initialized with the set of sources *S*. Then, A^⋆^ iteratively *expands* the explored state with lowest estimated cost by exploring all its neighbors, until it finds a target. Here, the cost for node *u* is estimated by the distance from source, called *g*(*u*), plus the estimate from the heuristic *h*(*u*).

### Heuristic Function

The heuristic function *h*(*u*) estimates the cost *h*^*^(*u*) of a shortest path in *G* from *u* to a target *t* ∈ *T*. Intuitively, a good heuristic correlates well with the distance from *u* to *t*.

To ensure that A^⋆^ indeed finds the shortest path, *h* should be *optimistic*:

#### Definition 1 (Optimistic heuristic)

*A heuristic h is optimistic if it provides a lower bound on the distance to the closest target: ∀u*.*h*(*u*) ≤ *h*^*^(*u*).

While any optimistic *h* ensures that A^⋆^ finds optimal alignments [6, Res. 3], the specific choice of *h* is critical for performance. In particular, decreasing the error *δ*(*u*) = *-h*^*^(*u*) *h*(*u*) can only improve the performance of A^⋆^[6, Res. 6]. Thus, a key contribution of ours is a domain-specific heuristic h.

### 3.2 AStarix: Instantiating A*

Algorithm 1 shows an unoptimized version of AStarix and its heuristic function. AStarix expects a reference graph (Line 1) and a query (Line 3) as input, and returns an optimal alignment (Line 7) by searching for a shortest path from *S* to *T* in the alignment graph 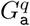. It is parameterized by hyper-parameters (*d* in Line 2, more in §4) and edit costs (implicitly provided).

The function Heuristic (Lines 8–11) computes a lower bound on the remaining cost of a best alignment: the minimum cost *h*(*u, s*) of aligning the *upcoming sequence s* (where |*s* | ≤*d*) starting from node *u*. Importantly, *s* is limited to the next *d*′ ≤ *d* letters of *q*, starting from query position *i*. Thus, computing *h*(*u, s*) is substantially cheaper than aligning all remaining letters of *q*.

To compute *h*(*u, s*) we leverage a simple branch-and-bound algorithm, provided in App. A.2. In the following, for convenience, we refer to the heuristic as *h* (which is parameterized by (*u, s*)) instead of Heuristic (which is parameterized by ⟨*u, i*⟩). Further, we say that *h* is optimistic if *h*(*u, s*) is a lower bound on the cost for aligning all remaining letters (i.e., *q*[*i* : |*q*|]) starting from node *u* (note that *s* is a prefix of *q*[*i* : |*q*|]).

#### Theorem 1. *h is optimistic*.

*Proof. h* only considers the next *d*′ letters of *q* instead of all remaining letters. Since all costs are non-negative, the theorem follows.

#### Benefit of A^⋆^Heuristic over Dijkstra

Fig. 2 shows the benefit of using our heuristic function compared to Dijkstra. Here, Dijkstra expands states based on their distance *g* from the origin nodes ⟨*u*, 0⟩ and ⟨*v*, 0⟩. Hence, depending on tie-breaking, Dijkstra may expand all states with *h* ≤1, as shown in Fig. 2. By contrast, A^⋆^ chooses the next state to expand by the sum of the distance from the origin *g* and the heuristic *h*, expanding only states with *g* +*h* ≤ 1.

**Fig. 2:**
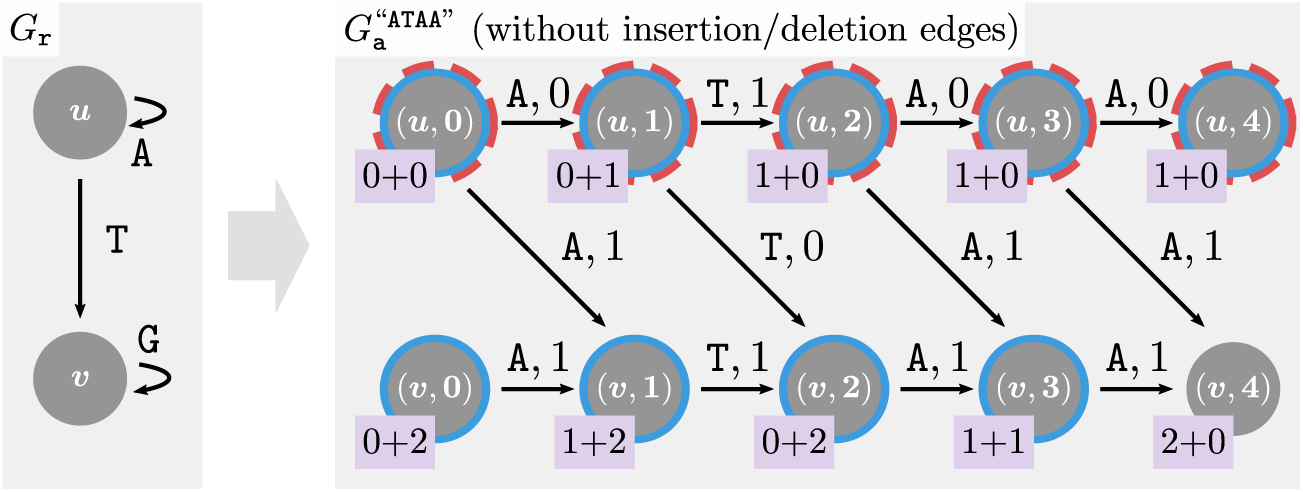
The benefit of using our heuristic over Dijkstra. Alignment graph 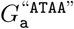 (right) is based on reference graph *G*_r_ (left), but omits insertion and deletion edges for simplicity. The pink boxes *g* + *h* indicate the distance from the sources *S* = {⟨*u*, 0⟩, ⟨*v*, 0⟩} (in *g*) and the cost of aligning the next *d* = 2 letters (in *h*). Dijkstra (resp. A^⋆^) expands states circled in blue (resp. dashed red).

#### Memoization

Recall that the return value of *h* in Line 8 only depends on *u* and the upcoming sequence *s* (which in turn depends on *i* and *d*). Thus, *h*(*u, s*) can be reused for different positions across different queries in 𝒪(1) time, if it was computed for a previous query.

## 4 AStarix Algorithm: Optimizations

We now discuss several optimizations we developed to speed up AStarix while preserving its optimality. These optimizations reduce preprocessing and alignment runtime as well as memory footprint (in particular for memoization).

### 4.1 Reducing Semi-global to Local Alignment Using a Trie

To find an optimal alignment, we generally need to consider all reference graph nodes *u* ∈*G*_r_ as possible starting nodes. Thus, optimal aligners PaSGAL [15] and BitParallel [27] brute-force through all possible starting nodes *u* ∈*G*_r_.

To more efficiently handle arbitrary starting positions for alignments, we extend the reference graph with a trie (referred to as *suffix tree* in [8]) to effectively align from all possible starting nodes *simultaneously*.

#### Single Starting State

In the trie approach, abstraction nodes are added to the graph, each of which corresponds to a set of nodes in *G*_r_ that correspond to the same prefix. In the following, we formalize this approach.

Concretely, we extend *G*_r_ by a *trie of depth D*, resulting in graph 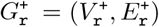. Our goal is that all paths in *G*_r_ that have length *D* and end in *v* ∈ *V*_r_ correspond to paths in 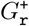 starting from a single source *ϵ* to 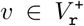, where *ϵ* represents the empty string. This correspondence ensures that it suffices to consider only paths in 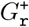 starting from the source *ϵ*. In particular, each alignment on 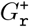 can be translated into an alignment on *G*_r_ (we omit this translation here).

Fig. 3 shows an example trie. To construct it, we first associate with every node *v* ∈ *V*_r_ the set 𝒮_*v*_ of its *D*-mers (orange boxes in Fig. 3): spells of paths ending in *v* and of length *D*. Our goal is then to use paths in the trie to spell these *D*-mers.

**Fig. 3:**
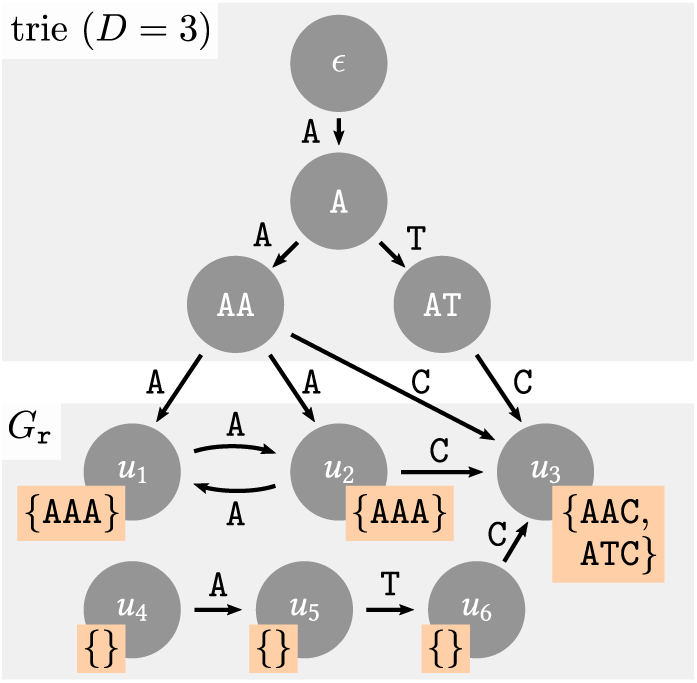
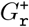 enables semi-global alignment by extending *G*_r_ with a trie.

Second, we construct the trie nodes from all prefixes of these *D*-mers:

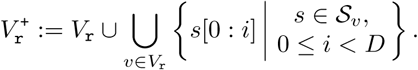

Third, we add edges within the trie, which ensure that paths from *ϵ* to any trie node *s* spell *s*. Formally, whenever 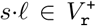, we add an edge (*s, s·l, l*) to 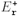, where “*·*” denotes string concatenation. Finally, we add edges between the trie and the reference graph, which ensure that any *D*-mer of any node *v* ∈*V*_r_ can be spelled by a walk from *E* to *v*. Formally, if *s·l* ∈ *𝒮*_*v*_, then 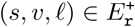.

Importantly, extending *G*_r_ to 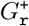 is compatible with the construction of the edit graph *G*_e_, the construction of the alignment graph and all other optimizations. In particular, when searching for a shortest path in the alignment graph constructed from 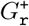, it suffices to only consider starting node ⟨*ϵ*, 0⟩.

#### Reducing Size of Trie

We can reduce the size of the trie by removing specific trie nodes. In particular, we iteratively remove each trie leaf node 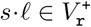 with a unique outgoing edge (*s l, v, l*′) to a reference graph node *v* ∈*V*_r_. To compensate for removing node *s l*, we introduce a new edge (*s, u, l*) to a node *u* ∈*V*_r_ with an edge (*u, v, l*′) (such a node must exist according to the construction of 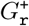). For example, in Fig. 3, we (i) remove node `AT` including its edges (`A`, `AT`, `T`) and (`AT`, *u*_3_, `C`), but (ii) introduce an edge (`A`, *u*_2_, *T*).

This optimization is lossless, as the *D*-mer *s · l · l*′ ∈ *S* _*v*_ can still be spelled by the path from *E* to *s*, extended by (*s, u, l*) and (*u, v, l*′).

### 4.2 Greedy Match Optimization

We also employ an optimization originally developed for computing the edit distance between two strings [31,1], but which has also been used in the context of string to graph alignment [8]. We omit the correctness proof of this optimization, which is already covered in [31], and only explain the intuition behind it.

Suppose there is only one outgoing edge *e* = (*u, v, f*) ∈ *E*_r_ from a node *u* ∈ *V*_r_. Suppose also that while aligning a query *q*, we explore state ⟨*u, i*⟩ for which the next query letter *q*[*i*] matches the label *f*. In this case, we do not need to consider the edit outgoing edges, because any edit at this point can be postponed without additional cost, as Δ_match_ ≤ min(Δ_subst_, Δ_ins_, Δ_del_). Thus, we can greedily explore state ⟨*v, i* + 1 ⟩, aligning *q*[*i* + 1] to *e* by using the edge (⟨*u, i* ⟩, ⟨*v, i* + 1 ⟩, *f*, Δ_match_) before continuing with the A^⋆^ search. We note that this optimization is only applicable when aligning in non-branching regions of the reference graph. In particular, it is not applicable for most trie nodes (§4.1).

### 4.3 Speeding Up Evaluation of Heuristic

In the following, we show how to reduce the runtime of evaluating the heuristic *h*(*u, s*), by introducing two separate optimizations that compose naturally.

#### Capping Cost

We cap *h*(*u, s*) at *c*, replacing it by *h*_*c*_(*u, s*) := min(*h*(*u, s*), *c*). To achieve this, we allow RecursiveAlign to ignore paths costing more than *c*. For large enough *c*, this speeds up computation without significantly decreasing the benefit of the heuristic, since nodes associated with a high heuristic value are typically not explored anyways. We investigate the effect of *c* in App. A.3.

##### Theorem 2.

*h*_*c*_ *is optimistic*.

*Proof*. We have *h*_*c*_(*u, s*) ≤ *h*(*u, s*) and that *h*(*u, s*) is optimistic (Theorem 1).□

#### Capping Depth

We reduce the number of nodes that need to be considered by *h*(*u, s*). To this end, we define a modified heuristic *h*_*d*_(*u, s*) that only considers nodes *R*_*u*_ ⊆*V*_e_ at distance at most *d* from *u* (cp. Line 2 in Algorithm 1): *R*_*u*_ :={*v* ∈ *V*_r_ | *∋* path *π* ∈ *G*_e_ from *u* to *v* with |*π*| ≤ *d* .}

If an alignment of *s* reaches the boundary of *R*_*u*_, defined as

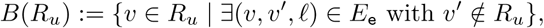

it is allowed to only spell a prefix of *s*, and the remaining unaligned letters of *s* are considered aligned with zero cost:

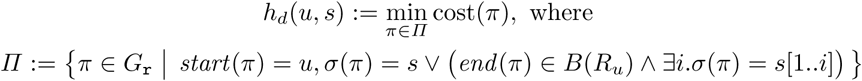

##### Theorem 3.

*h*_*d*_ *is optimistic*.

*Proof*. It suffices to show *h*_*d*_(*u, s*) ≤*h*(*u, s*) since *h*(*u, s*) is optimistic. In the case where all of *s* is aligned, *h*_*d*_(*u, s*) = *h*(*u, s*). Otherwise, the unaligned letters of *s* are not penalized, so *h*_*d*_(*u, s*) ≤ *h*(*u, s*). □

#### 4.4 Partitioning Nodes into Equivalence Classes

We have shown in §3.2 how to reuse an already computed *h*(*u, s*) for repeating *s* across different queries and query positions. In the following, we additionally aim to reuse *h*(*u, s*) across different nodes *u*, so that *h*(*u, s*) does not need to be computed for all nodes *u*. Intuitively, we want to assign two nodes *u* and *v* to the same equivalence class when the *graph region* considered by *h*(*u, s*) is equivalent to the graph region considered by *h*(*v, s*), up to renaming of nodes.

Thus, *h*(*u, s*) = *h*(*v, s*) if *u* and *v* are from the same equivalence class. Therefore, we can (arbitrarily) choose a representative node *r* ∈ *V*_r_ for every equivalence class, and evaluate *h*(*r, s*) instead of *h*(*u, s*), where *r* is the representative of the equivalence class of *u*. To look up representative nodes in 𝒪(1), we define a helper array *repr* with *repr* [*u*] = *r*.

#### Identifying Equivalence Classes

To identify the nodes belonging to the same equivalence class, we assume the optimization from §4.3, i.e., that our heuristic only considers nodes up to a distance *d* from *u*. Moreover, for performance reasons, our implementation detects only the equivalence classes of nodes *u* with a single outgoing path of length at least *d*. In this case, *u* and *u′* are in the same equivalence class if their outgoing paths spell the same sequence. In contrast, we leave nodes with forking paths in separate equivalence classes.

Note that for smaller *d*, the number of equivalence classes gets smaller, the reuse of the heuristic gets higher, and the memoization table has a lower memory footprint. At the same time, however, the heuristic *h*_*d*_(*u, s*) is less informative.

## 5 Evaluation

In this section we present a thorough experimental evaluation^3^ of AStarix on simulated Illumina reads. Our evaluation demonstrates that:

1. AStarix is faster than Dijkstra because the heuristic reduces the number of explored states by an order of magnitude.
2. The runtime of AStarix scales better than state-of-the-art optimal aligners with increasing graph size, on a variety of reference graphs.

### 5.1 Implementation of AStarix and Dijkstra

Our AStarix implementation uses an adjacency list graph data structure to represent the reference and the trie in a unified way, representing each letter by a separate edge object. To represent the reverse complementary walks in *G*_r_, the vertices are doubled, connected in the opposite direction, and labeled with complementary nucleotides (`A` ↔ `T`, `C` ↔ `G`). We do not limit the number of memoized heuristic function values (§3.2), but note we could do so by resetting the memoization table periodically. Our implementation of Dijkstra reuses the same AStarix codebase except the use of a heuristic function (i.e., with *h* ≡0).

We apply all described optimizations to AStarix and Dijkstra, except §4.3 and §4.4 which are applicable only to AStarix.

While the optimality of AStarix is not affected by its parameters, its performance is (see App. A.3 for analysis). To compare with other aligners, we use values *d* = 5, *c* = 5, *D* = ⌊log*Σ* |*G*_r_|⌋.

### 5.2 Compared Aligners: PaSGAL and BitParallel

We compare the performance of AStarix to that of two state-of-the-art optimal aligners: PaSGAL and BitParallel, with their default parameters. We do not compare to the exact aligner of VG as (i) its optimal alignment is intended for testing purposes only, (ii) it does not provide an interface for aligning a set of reads, and (iii) it has been consistently outperformed by PaSGAL [15].

PaSGAL is compiled with AVX2 SIMD support. The resulting alignments are not expected to match exactly between the local aligner PaSGAL and the semi-global aligners (AStarix and BitParallel) as they solve different tasks with different edit costs. Nevertheless, in analogy with the evaluations of PaS-GAL [15], it is still meaningful to compare performance, assuming that the dynamic programming approach of PaSGAL can be adapted to semi-global alignment with similar performance.

Both BitParallel and PaSGAL reach their worst-case runtime complexity independent of the edit costs Δ = (Δ_match_, Δ_subst_, Δ_ins_, Δ_del_). PaSGAL is evaluated using its default costs Δ = (*-*1, 1, 1, 1) and BitParallel is evaluated using the only supported costs Δ = (0, 1, 1, 1).

### 5.3 Setting

All evaluations were executed singled-threaded on an Intel Core i7-6700 CPU running at 3.40GHz.

#### Reference Graphs and Reads

We designed three experiments utilizing three different reference graphs (in Table 1). The first is a linear graph without variation based on the *E. coli* reference genome (strain: K-12 substr. MG1655, ASM584v2 [13]). The other two are variation graphs taken from the PaSGAL evaluations [15]: they are based on the Leukocyte Receptor Complex (LRC, with 1 099 856 nodes and 1 144 498 edges), and the Major Histocompatibility Complex (MHC1, with 5 138 362 nodes and 5 318 019 edges). We note that we do not evaluate on de Brujin graphs, since PaSGAL does not support cyclic graphs.

**Table 1:** Performance of optimal aligners for different reference graphs.

| Genome graph | Size | Runtime and Memory |  |  |  |
| --- | --- | --- | --- | --- | --- |
|  |  | ASTARIX | DIJKSTRA | PASGAL | BITPARALLEL |
| *E. coli* (linear) | ~4.7 Mbp | 33 sec<br>0.66 GB | 73 sec<br>0.66 GB | 3 272 sec<br>0.55 GB | 4 906 sec<br>0.43 GB |
| LCR (variant) | ~1 Mbp | 437 sec<br>1.12 GB | 940 sec<br>1.09 GB | 1 614 sec<br>0.30 GB | SegFault |
| MHC1 (variant) | ~5 Mbp | 1 282 sec<br>4.35 GB | 1 588 sec<br>1.21 GB | >7 200 sec<br>0.87 GB | SegFault |

**Table 2:**
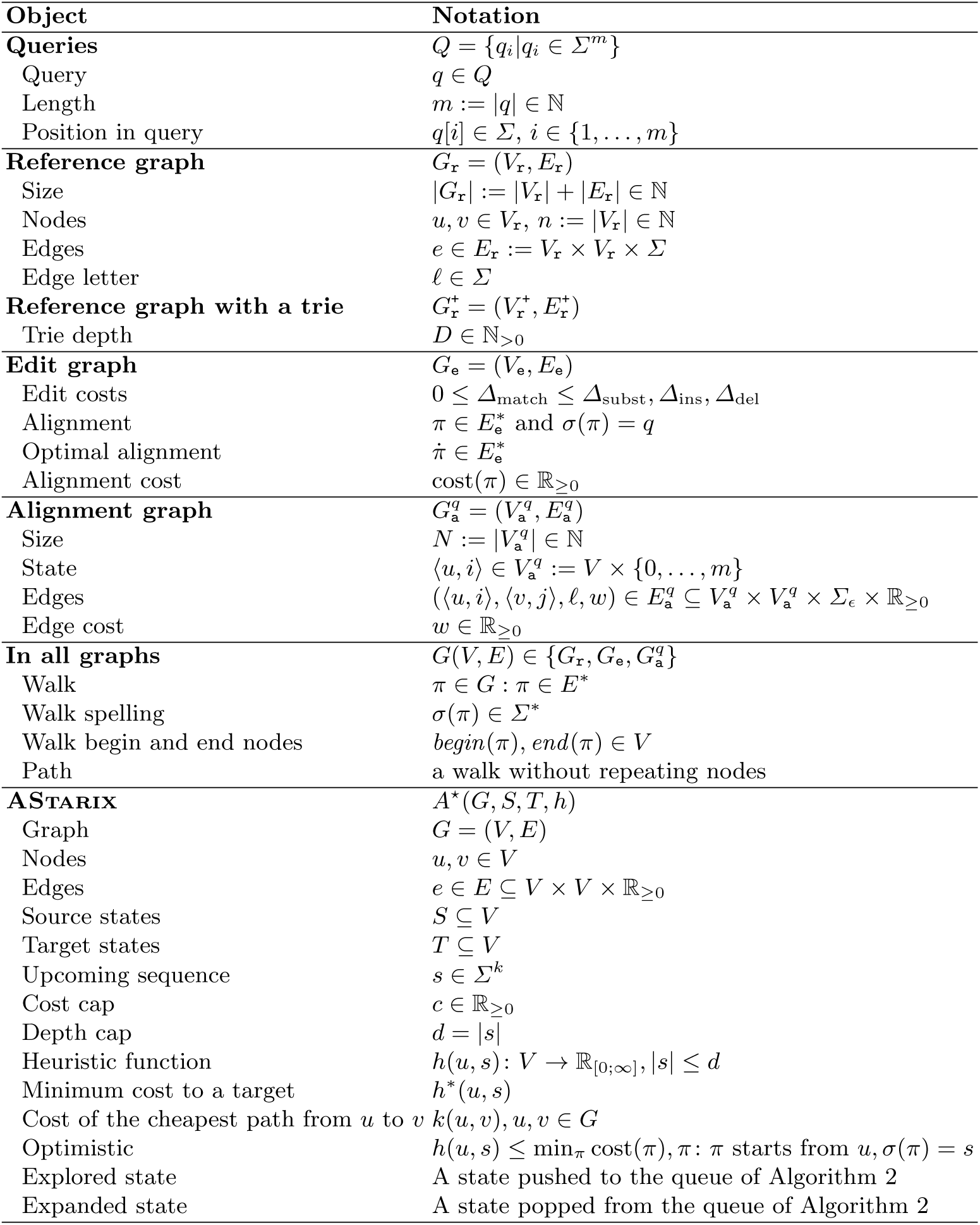
Notational conventions.

For the *E. coli* dataset we used the ART tool [14] to simulate an Illumina single-end read set with 10 000 reads of length 100. For the LCR and MHC1 datasets, we sampled 20 000 single-end reads of length 100 from the already generated sets in [15] using the Mason2 [12] simulator.

For Dijkstra and AStarix, the runtime complexity depends not only on the data size, but also on the data content, including edit costs. More accurate heuristics lead to better A^⋆^ performance [26], which is why we evaluate AStarix with costs corresponding more closely to Illumina error profiles: Δ = (0, 1, 5, 5).

#### Metrics

As all aligners evaluated in this work are provably optimal, we are mostly interested in their performance. To study the end-to-end performance of the optimal aligners, we use the Snakemake [20] pipeline framework to measure the execution time of every aligner (including the time spent on reading and indexing the reference graph input and outputting the resulting alignments). We note that the alignment phase dominates for all tools and experiments.

To judge the potential of heuristic functions, we measure not only the runtime but also the number of states explored by AStarix and Dijkstra. This number reflects the quality of the heuristic function rather than the speed of computation of the heuristic, the implementation and the system parameters.

### 5.4 Comparison of Optimal Aligners

#### Different Reference Graphs

Table 1 shows the performance of optimal aligners across various references. On all references, AStarix is consistently faster than Dijkstra, which is consistently faster than PaSGAL and BitParallel. The memory usage of Dijkstra is within a factor of 3 compared to PaSGAL and BitParallel. Due to the heuristic memoization, the memory usage of AStarix can grow several times compared to Dijkstra.

#### Scaling with Reference Graph Size

Fig. 4 compares the performance of existing optimal aligners. BitParallel and PaSGAL always explore all states, thus their average-case reaches the worst-case complexity of 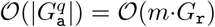. Due to the trie indexing, the runtime of AStarix and Dijkstra scales in the reference size with a polynomial of power around 0.2 versus the expected linear dependency of BitParallel and PaSGAL.

**Fig. 4:**
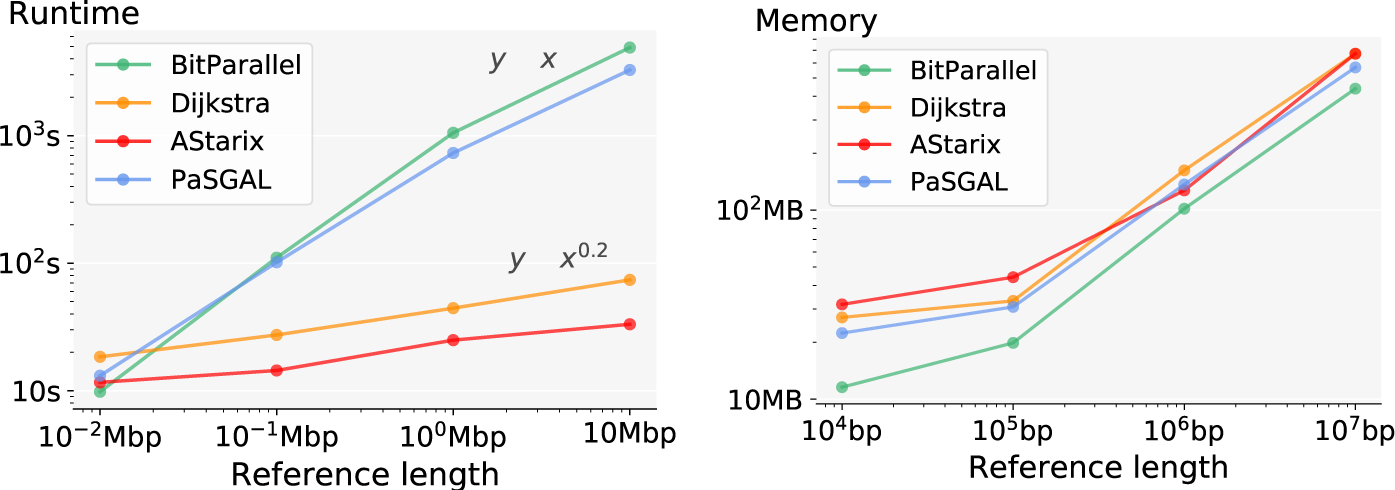
Comparison of overall runtime and memory usage of optimal aligners with increasing prefixes of E. coli as references.

The heuristic function of AStarix demonstrates a 2-fold speed-up over Dijkstra. This is possible due to the highly branching trie structure, which allows skipping the explicit exploration for the majority of starting nodes.

### 5.5 A^⋆^ Speedup

To measure the speedup caused by the heuristic function, we compare the number of not only the expanded, but also of explored states (the latter number is never smaller, see §3.1 and the example in Fig. 2) between AStarix and Dijkstra on the MHC1 dataset.

Fig. 5 demonstrates the benefit of the heuristic function in terms of both alignment time and number of explored states. Most importantly, Astarix scales much better with increasing number of errors in the read, compared to Dijkstra. More specifically, the number of states explored by Dijkstra, as a function of alignment cost, grows exponentially with a base of around 10, whereas the base for AStarix is around 3 (the empirical complexity is estimated as a best exponential fit *exploredStates* ∼*a ·score*^*b*^).

**Fig. 5:**
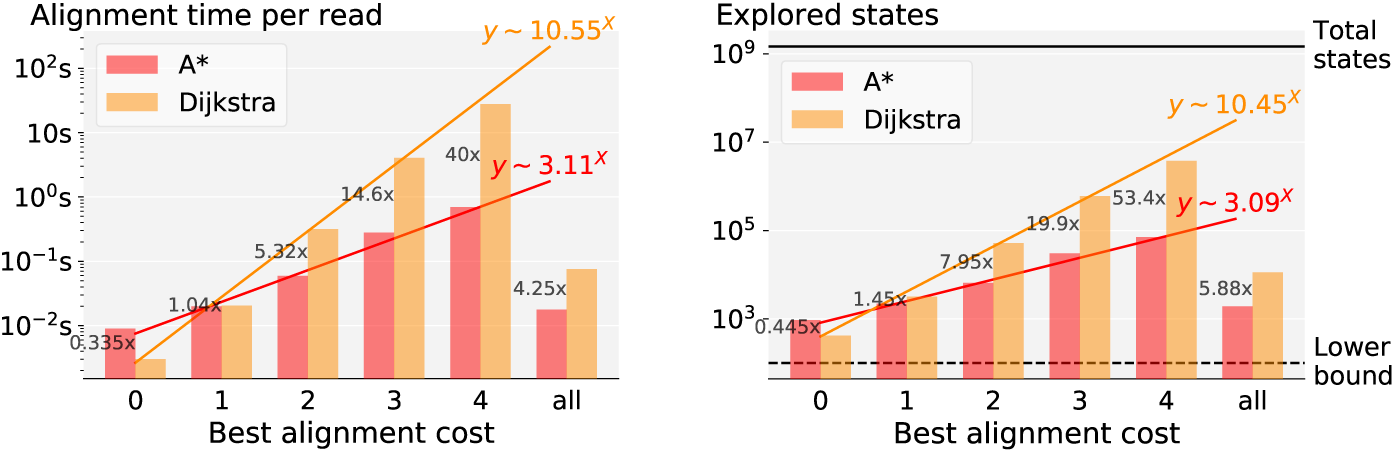
Comparison of A^⋆^ and Dijkstra in terms of mean alignment runtime per read and mean explored states depending on the best alignment cost on MHC1.

**Fig. 6:**
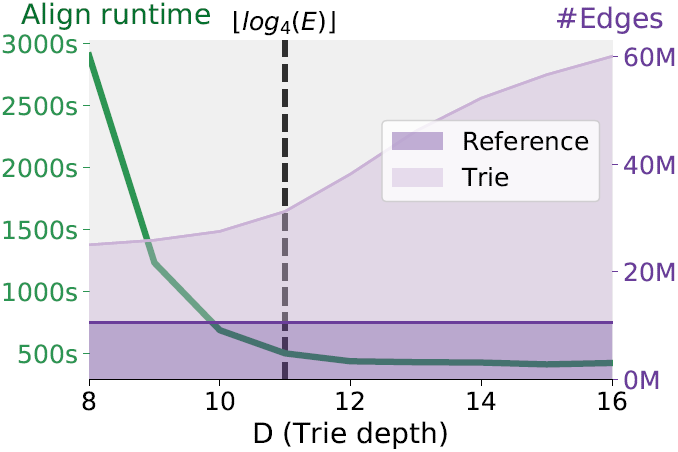
Effect of *D* on performance of AStarix (MHC1 experiment). The dashed line shows our choice of *D*.

**Fig. 7:**
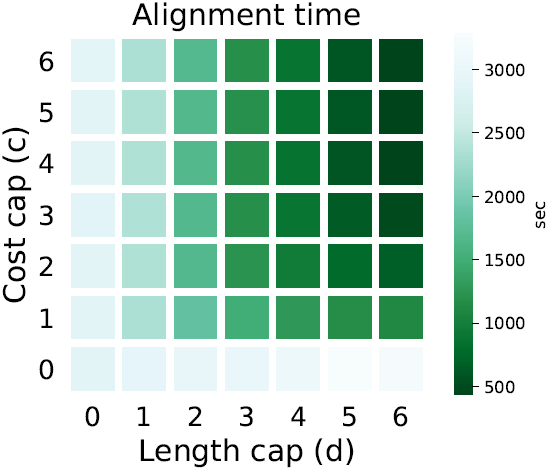
Runtime of AStarix depending on *d* and *c* (MHC1 experiment).

The horizontal black line in Fig. 5 denotes the total number of states |*G*_r_| · |*q* |, which is always explored by BitParallel and PaSGAL. On the other hand, any aligner must explore at least *m* = |*q*| states, which we show as a horizontal dashed line. This lower bound is determined by the fact that at least the states on a best alignment need to be explored.

## 6 Conclusion

We presented AStarix, an A^⋆^ algorithm to find optimal alignments, based on a domain-specific heuristic and enhanced by multiple algorithmic optimizations. Importantly, our approach allows for both cyclic and acyclic graphs including variation and de Bruijn graphs.

We demonstrated that AStarix scales exponentially better than Dijkstra with increasing (but small) number of errors in the reads. Moreover, for short reads, both AStarix and Dijkstra scale better and outperform current state-of-the-art optimal aligners with increasing genome graph size. Nevertheless, scaling optimal alignment of long reads on big graphs remains an open problem.

We expect that AStarix can be scaled further, to both (i) bigger graphs and (ii) longer and noisier reads. Scaling AStarix may require a combination of (i) the development of more clever heuristic functions (by leveraging existing work on A^⋆^ and edit distance) and (ii) algorithmic optimizations. We note that if desired, a (sub-optimal) seeding step could speed up AStarix by pre-filtering the starting positions, analogously to other practical aligners.

## A Appendix

### A.1 Generic Algorithms: A^⋆^ and Dijkstra

Algorithm 2 shows a generic implementation of the A^⋆^ algorithm, roughly following [6]. We do not implement the reconstruction of the best alignment in order to simplify the presentation. The procedure BacktrackPath traces the best alignment back to the *source*, based on remembered edges used to optimize *f* for each alignment state. Algorithm 2 also shows a simple implementation of Dijkstra in terms of A^⋆^.

#### Algorithm 2 A^⋆^ algorithm (generalizes Dijkstra)

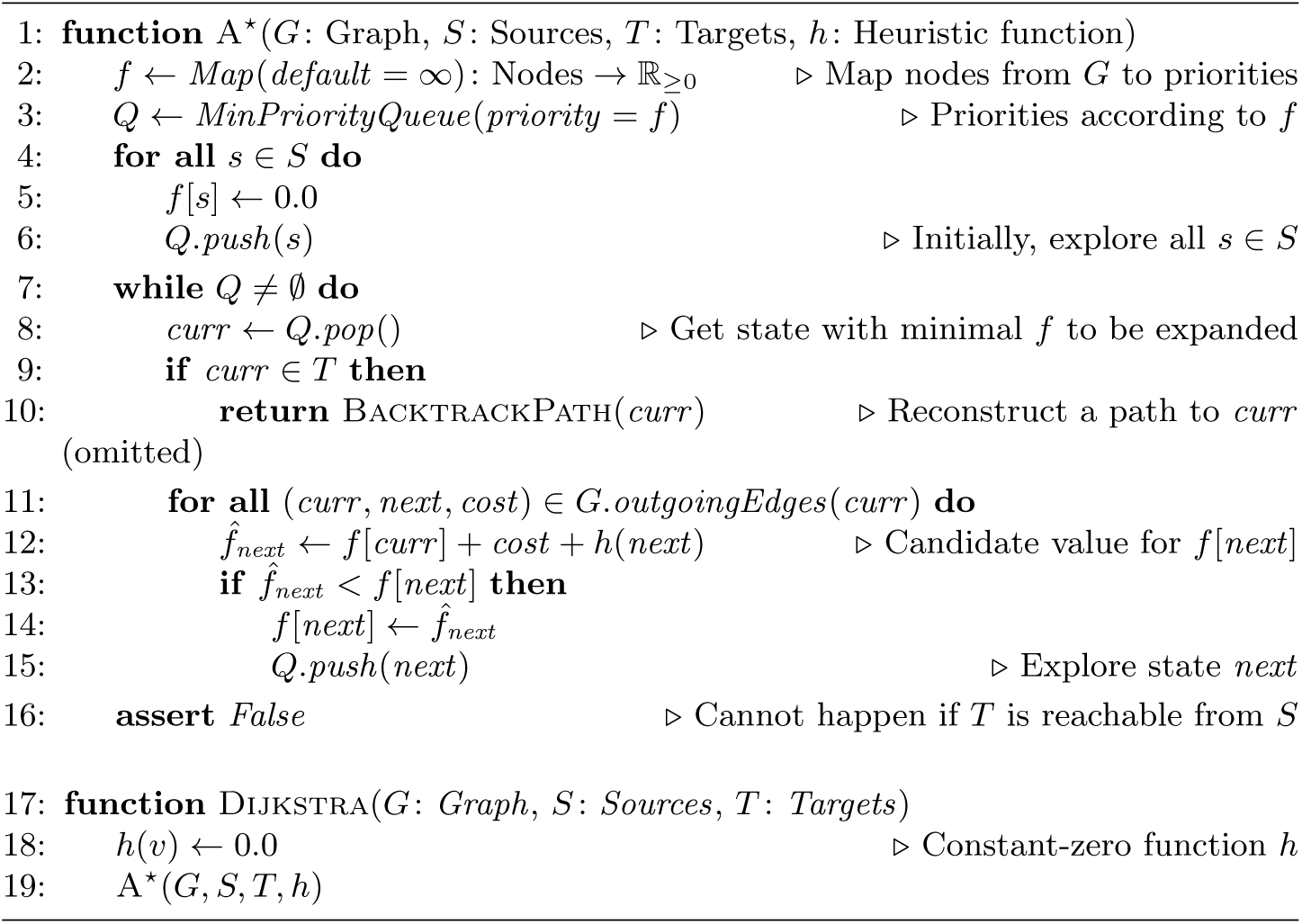

#### Algorithm 3 Recursive alignment used by Heuristic in Algorithm 1.

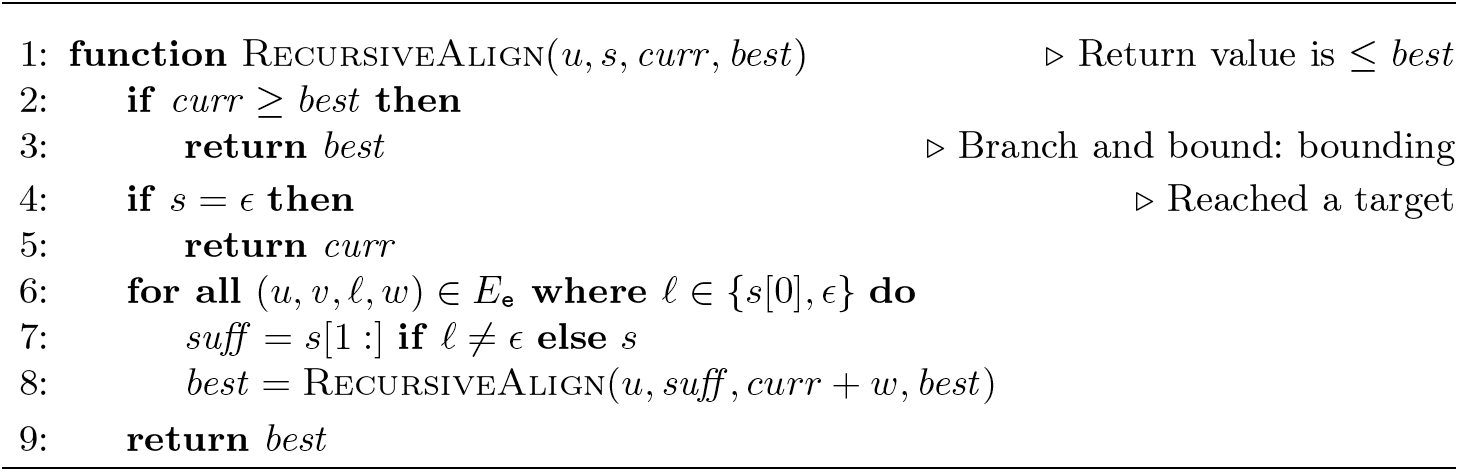

### A.2 Recursive Alignment Algorithm

Algorithm 3 shows our implementation of RecursiveAlign, used in Algorithm 1 to evaluate *h*. RecursiveAlign is a simple branch-and-bound algorithm that recursively looks for the cheapest alignment of *s* starting from *u*, and does not follow paths whose cost exceeds *best*, the best path found so far.

### A.3 Parameter Estimation

We now evaluate the influence of different parameter choices (*c, d, D*) on runtime and memory usage.

Fig. 6 demonstrates the benefit of using a trie with the size reduction optimization (end of §4.1): increasing the trie depth *D* speeds up aligning but requires more memory. Selecting the trie depth based on the graph size *D* = ⌊log*Σ* |*G*_r_|⌋ provides a reasonable trade-off between alignment time and memory.

Fig. 7 shows the joint effect of *c* and *d*. It demonstrates that having a long reach (*d*) that covers at least some errors (*c >* 0) is a reasonable strategy for choosing *d* and *c*.

### A.4 Versions, commands, parameters for running all evaluated approaches

In the following, we provide details on how we executed the approaches discussed in §5:

#### PaSGAL

Obtained from https://github.com/ParBLiSS/PaSGAL (Commit `50ad80c`)

Command PaSGAL -q reads.fq -r graph.vg -m vg -o output -t 1

#### BitParallel

Obtained from https://github.com/maickrau/GraphAligner/tree/WabiExperiments (Commit `241565c`)

Command `Aligner -f reads.fq -g graph.gfa >output`

#### AStarix

Obtained from https://github.com/eth-sri/astarix/tree/recomb2020

Command `astarix align-optimal -f reads.fq -g graph.gfa >output`

#### Dijkstra

Obtained from https://github.com/eth-sri/astarix/tree/recomb2020

Command `astarix align-optimal -f reads.fq -g graph.gfa –a dijkstra >output`

### A.5 Notations

Table 2 summarizes the notational conventions used in this work.

## Footnotes

1 We refer as BitParallel to to the bit-parallel DP algorithm implemented in GraphAligner tool [27].

2 The appendix with algorithms and evalution details is included in the full version of this paper: https://www.biorxiv.org/content/10.1101/2020.01.22.915496v1

3 https://github.com/eth-sri/astarix/tree/RECOMB2020_experiments

